# Keratin 7 protein presence in stool is indicative of active pediatric-onset inflammatory bowel disease

**DOI:** 10.64898/2026.04.21.719629

**Authors:** Maria A. Ilomäki, Ekta Kotharkar, Jannika Rovapalo, Noora Lehtonen, Anne Nikkonen, Rebecka Ventin-Holmberg, Johannes Merilahti, Otto Kauko, Kaija-Leena Kolho, Lauri Polari, Diana M. Toivola

## Abstract

**Background:** Inflammatory bowel disease (IBD) is associated with early structural changes in intestinal epithelial cells; however, the associated molecular alterations remain incompletely understood. The cytoskeletal protein keratin (K) 7 was recently found to be focally expressed in the colonic epithelium in IBD, while absent in the healthy colon. Here, we investigated the applicability of K7 as a noninvasive stool biomarker for pediatric IBD.

**Methods:** In this case-control study including adolescent patients with IBD (n=27) and healthy controls (n=15), stool lysates were analyzed by proteomics, immunoassay and qPCR to determine K7 protein and mRNA content, respectively. Additionally, stool mRNA levels of the simple epithelial keratins, K8, K18, K19 and K20, were measured.

**Results:** Stool proteomic analysis identified focal K7 and K19 in IBD samples. Additionally, 23 differentially abundant proteins, of which 18 were higher in IBD, were identified and Gene Ontology enrichment analysis highlighted immune and inflammatory pathways. K7 specific immunoassay detected fecal K7 protein in all patients with active IBD, including both ulcerative colitis and Crohn’s disease, while K7 was near or below the detection limit in controls and IBD patients in remission (area under ROC curve=0.88, p<0.0001). While *KRT7* mRNA levels were below the detection limit, *KRT8* and *KRT18* transcripts were elevated in IBD samples compared to controls (p<0.05).

**Conclusions:** K7 protein is elevated in IBD patient stool, reflecting intestinal de novo expression and increased epithelial cell exfoliation. Fecal K7 may provide a novel, noninvasive marker for IBD diagnosis and monitoring.

## Introduction

Inflammatory bowel disease (IBD) has been linked to a disruption of the intestinal barrier homeostasis maintained between the immune system, the intestinal microbiota and epithelial cells [1]. Defects in the intestinal barrier have been suggested as an early, preclinical step in the pathogenesis of both ulcerative colitis (UC) and Crohn’s disease (CD) [2–4]. Active IBD is associated with epithelial pathologies including mucin producing cell depletion, abnormal crypt architecture, and erosion [5,6]. IBD-induced changes in epithelial cell fate, transcriptome and function have been reported earlier [7,8], but the involvement of cytoskeletal composition in barrier breakdown is not fully understood.

Keratins (K) are a group of intermediate filament forming proteins and represent major cytoskeletal components of epithelial cells. In the intestinal epithelia the primary keratins include type II K8, type I K18, K19 and K20 [9]. They maintain the homeostasis of the epithelial cell layer through both cell-cell and cell-basal layer connections [10]. While keratin expression changes in colorectal cancer have been commonly utilized in pathology, changes in other colonic diseases have been less of a focus [11]. Recently, we found that K7 is focally expressed *de novo* in colonocytes of patients with IBD, both on a protein and mRNA level [12,13]. K7 is normally expressed in ductal and glandular cells but not in the healthy colonic epithelium [14]. Additionally, earlier studies utilizing both human intestinal biopsies and colitis mouse models have observed inflammation-associated changes in the expression of intestinal keratins [15–18].

Previously we showed that fecal keratin mRNA content mirrors colonic epithelial keratin expression and that fecal K19 mRNA is elevated in stool during intestinal epithelial damage in mice [19]. Furthermore, clinical studies have demonstrated the feasibility of using exfoliated epithelial cells in stool for noninvasive detection of epithelial cell gene expression [20,21], and the potential of fecal K8 as a biomarker in necrotizing enterocolitis in newborns [22]. Along with our discovery that K7 is neo- expressed specifically in IBD colon, these findings encourage diagnostic applications for keratins.

Currently the most widely utilized IBD biomarker, neutrophil produced fecal calprotectin (FC), is not IBD-specific, as it is sporadically elevated in various common gut diseases [23,24]. Timely use of colonoscopy for diagnosis and monitoring of IBD remains the standard, however, the lack of specific biomarkers for noninvasive predicting and monitoring of the disease remains a clinical gap [25]. This study assessed fecal levels and changes in K7 and other intestinal epithelial proteins in adolescent patients with IBD compared to non-IBD controls, using proteomic analysis, immunoassay, and qPCR.

## Materials and Methods

### Study Design and Subjects

The cohort consisted of 42 donors (Table 1) recruited between 2011 and 2021 at the Children’s Hospital, Helsinki University and HUS, Helsinki, Finland, when undergoing colonoscopy or during their follow-up. Control subjects (CTRL) comprised children who remained without an IBD diagnosis after colonoscopy (n = 11) and healthy children without gastrointestinal conditions (n = 4).

**Table 1.** Clinical chart of patient characteristics.

| Characteristics | n (%) or Median (min-max) |  |
| --- | --- | --- |
|  | Control | IBD |
| No. of patients | 15 (35.7) | 27 (64.3) |
| Ulcerative colitis (UC) |  | 12 (28.6) |
| Crohn's disease (CD) |  | 11 (40.7) |
| IBD unclassified (IBDU) |  | 4 (9.5) |
| Age at sampling, years | 12.8 (9.7-16.8) | 15.4 (9.5-18.7) |
| Age at diagnosis, years |  | 12.1 (4.9-16.2) |
| Female | 6 (14.3) | 10 (23.8) |
| FC at sampling (µg/g) | 10 (<5-40) | 605 (11-5979) |
| Medication at sampling |  |  |
| Glucocorticoid |  | 8 (19.0) |
| 5-aminosalicylic acid |  | 11 (26.2) |
| Azathioprine |  | 3 (7.1) |
| Methotrexate |  | 4 (9.5) |
| Biological medication |  | 11 (26.2) |
| Antibiotics |  | 2 (4.8) |

IBD = inflammatory bowel disease, FC = fecal calprotectin

All patients or guardians signed an informed consent form to participate in a study on pediatric-onset IBD, approved by the Ethics Committee of the Hospital District of Helsinki and Uusimaa (371/E7/2004), (97/13/03/03/2011), (351/13/03/03/2012), (198/13/03/01/2014), and (HUS/1170/2020).

### Fecal Sample Handling

Samples consisted of 29 IBD and 15 CTRL fecal specimens, with two UC samples representing longitudinal follow-up specimens. Among the IBD samples, 6 were defined as in remission (FC levels <100 µg/g) and 22 as active disease (FC >100 µg/g). FC levels were measured following previous protocol [26]. Samples were collected at home and immediately frozen at −20 °C or stored for 1–3 days at +4 °C or room temperature before storage at −70 °C. Three samples were thawed once before analysis. Samples were aliquoted on dry ice for subsequent analyses (Figure 1).

**Figure 1.**
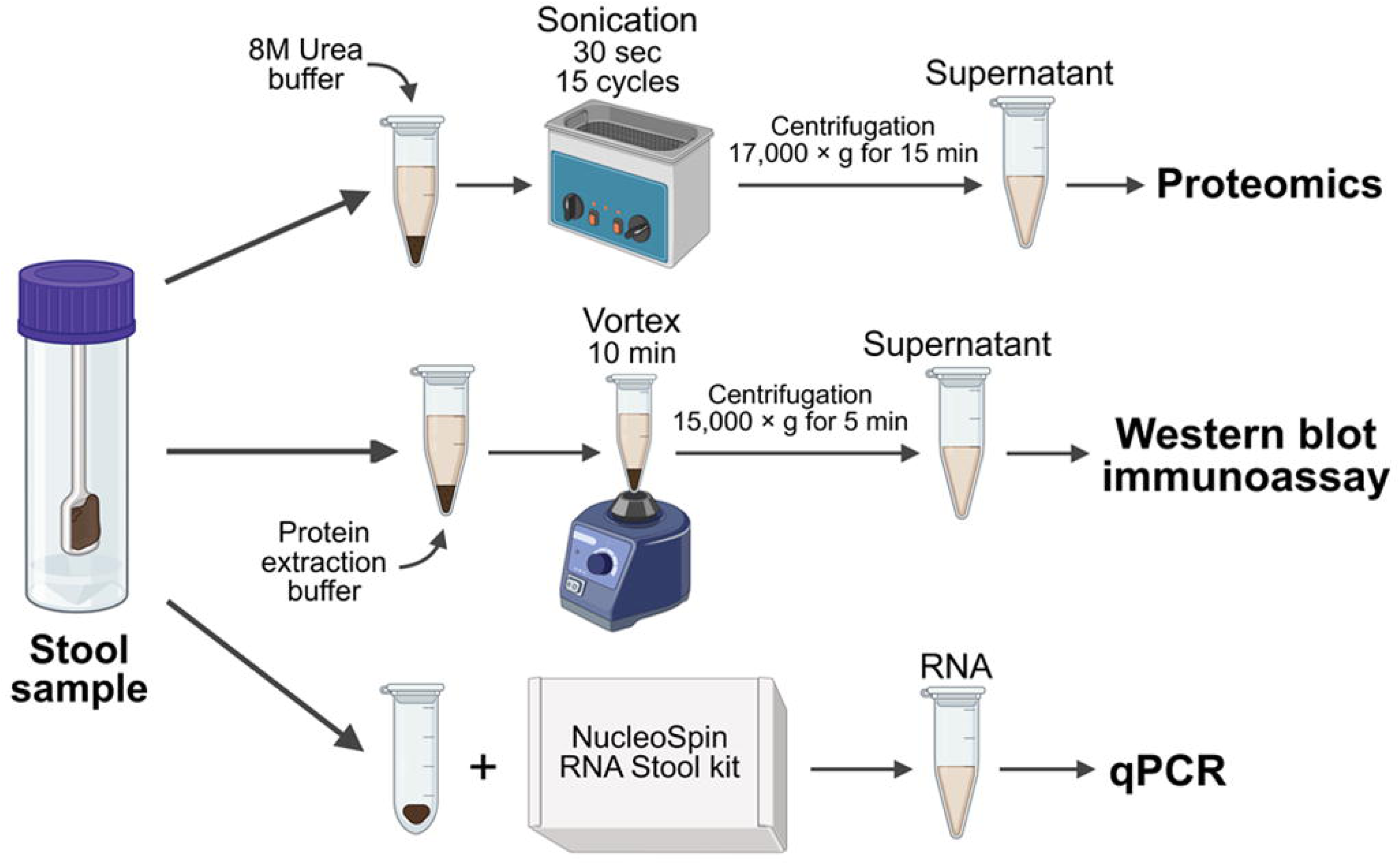
Simplified flow chart on the methods of protein and RNA extraction from stool.

### Stool Proteomics

Proteomics was conducted on 14 IBD (5 UC, 6 CD, 3 IBDU) and 9 CTRL samples. Among the IBD group, 6 patients were in remission and 8 had active disease. Samples were anonymized and randomized for processing. Aliquots were denatured in urea buffer (8 M urea, 50 mM Tris-HCl (pH 8.0), cOmplete™ Mini EDTA-free protease inhibitor cocktail (Roche, cat. no. 11836170001)), then prepared at +4 °C according to Figure 1, followed by 10 mM tris(2-carboxyethyl) phosphine, 40 mM chloroacetamide (30 minutes, room temperature) and overnight digestion using Trypsin/Lys-C mix (Promega, Madison, WI). Peptides were acidified, and 500 ng loaded onto Evotips for Evosep One system (Evosep Biosystems, Odense, Denmark) chromatographic separation and data-independent acquisition (DIA) – Parallel Accumulation Serial Fragmentation (PASEF) LC–MS/MS analysis using a timsTOF FLEX mass spectrometer (Bruker, Bremen, Germany).

Separation was performed on a Evosep Performance column (150 μm × 15 cm, 1.5 μm C18 beads) using water with 0.1% formic acid and 0.1% formic acid in 99.9% acetonitrile. The DIA–PASEF method was applied using a 44-minute gradient. Data was acquired using Compass 2025b software (Bruker) across an m/z range of 301–1100. Each run comprised 25 DIA–PASEF scans with a ramp period of 100 ms and two ion mobility windows (0.6-1.3 V·cm⁻²) and variable isolation widths.

### Mass Spectrometry Data Processing and Quantification

Raw data was processed using Spectronaut v20.1 (Biognosys AG, Schlieren, Switzerland) and DirectDIA workflow. Protein searches were performed against the SwissProt Homo sapiens database (release 2025_01) according to previous parameters [27], supplemented with the Universal Protein Contaminant database [28]. The false discovery rate was set at 1% at both precursor and protein levels. Quantification was performed at the MS2 level using fragment ion under the curve (AUC) with local normalization based on a retention time–dependent regression model [28,29].

### Differential Abundance and Gene Ontology Enrichment Analysis

Principal component analysis (PCA) and differential abundance analysis were conducted in Spectronaut (v20.2) using normalized protein quantities. Gene Ontology (GO) enrichment was conducted using protein lists derived from differential abundance analysis. The enrichGO function from the clusterProfiler package (v4.16.0) was used for GO Biological Process (BP) overrepresentation analysis. Gene symbols were converted to Entrez IDs using the bitr() function and the org.Hs.eg.db annotation database (v3.17.0). GO enrichment was conducted using the BP ontology in clusterProfiler with the default genome-level background annotation and significance thresholds of p<0.05 and Benjamini–Hochberg adjusted q<0.2.. Enriched GO terms were visualized using clusterProfiler’s dotplot() function.

### Western Blot Immunoassay

Samples were diluted 150 mg stool/1 ml calprotectin extraction buffer (Actim, Espoo, Finland) with 1× complete protease inhibitor cocktail (Roche, Basel, Switzerland) and 1 mM phenylmethylsulfonyl fluoride (Sigma-Aldrich, St. Louis, MO), then prepared at +4 °C according to Figure 1. K7-negative control lysates were made from yeast using the same protocol.

Sample lysates were mixed 1:1 with Laemmli sample buffer prepared according to previous protocol [19], and separated on 10 % SDS-polyacrylamide gels together with Precision Plus Protein Dual Color Standards (Bio-Rad, Hercules, CA). Yeast lysate and recombinant 6xHis-tagged K7 (Cusabio, Wuhan, China) were loaded as a negative and positive control, respectively. Western blot analysis was performed following previous protocol [19], using rabbit anti-K7 D1E4 diluted 1:1000 (Cell Signaling Technology, Danvers, MA) as the primary antibody and anti-rabbit HRP diluted 1:10,000 (Promega) as the secondary antibody. Signal was developed using Western Lightning Plus-ECL (Revvity, Waltham, MA, USA). Bands were quantified by two independent scientists using ImageJ software (National Institutes of Health, Bethesda, MD).

### Gene Expression Analysis

RNA was extracted from the stool samples using the NucleoSpin RNA Stool kit (Macherey Nagel, Düren, Germany) according to the manufacturer’s protocol (Figure 1). Total RNA was determined using Nanodrop 2000 Spectrophotometer (Thermo Fisher Scientific). RNA samples were reverse transcribed to cDNA following the manufacturer’s protocol (Promega). Genes of interest were quantitated using QuantStudio 3 real-time PCR system (Applied Biosystems, Waltham, MA) with TaqMan Universal Master Mix II, no UNG, and TaqMan gene expression Assay (Supplemental Table 1), following the manufacturer’s qPCR protocol. Results are presented as qPCR cycle threshold (Ct) values.

### Statistical Analysis and Image processing

Proteomics differential abundance analysis was conducted using the Spectronaut software with unpaired Student’s t-test on log₂ transformed protein quantities, with significance thresholds set at p ≤ 0.05, |log₂ fold change| ≥ 0.58 and the q-value set to 1. The venn diagram was built in R (version 4.5.1) using the VennDiagram package (v1.7.3). Proteins were considered present in a group if detected in at least one sample within that group. Volcano plots and GO enrichment figures were generated in R (version 4.5.1). For the proteomics, Westen blot and qPCR data, graphs were generated and statistically analysed using GraphPad Prism version V9.0 (GraphPad Software Inc., San Diego, CA). Differences between two groups and three groups or more were determined using unpaired t-test and Kruskal-Wallis test, respectively. Correlation between parameters was determined using Pearsons. To evaluate the clinical performance of keratins as IBD biomarkers, receiver operating characteristics (ROC) curves were generated and AUC with a 95% confidence interval was calculated.

Figure 1 was created using BioRender.com (Science Suite Inc. Toronto, Canada), the rest of the figures generated with Adobe Illustrator 2025 (Adobe, Inc., San Jose, CA).

## Results

### Stool Proteomics Analysis Identifies IBD-associated Protein Profiles and Detects Intestinal Keratins

PCA of the stool proteome revealed separation between IBD and CTRL samples along PC1 (21.4%) and PC2 (18.2%) (Figure 2A). When the IBD group was divided into active and remission states, active IBD samples were distinctly separated, whereas remission samples overlapped with the CTRL samples (Figure 2B). Using DIA-based proteomics, 950 human proteins were identified (Supplemental Table 2), with their shared and unique distributions displayed in the Venn diagram (Figure 2C). 494 proteins were shared among CTRL, active IBD, and remission samples, representing a common stool proteome. Moreover, 292 proteins were detected only in active IBD, while 3 were unique for IBD in remission and 7 for CTRL samples.

**Figure 2.**
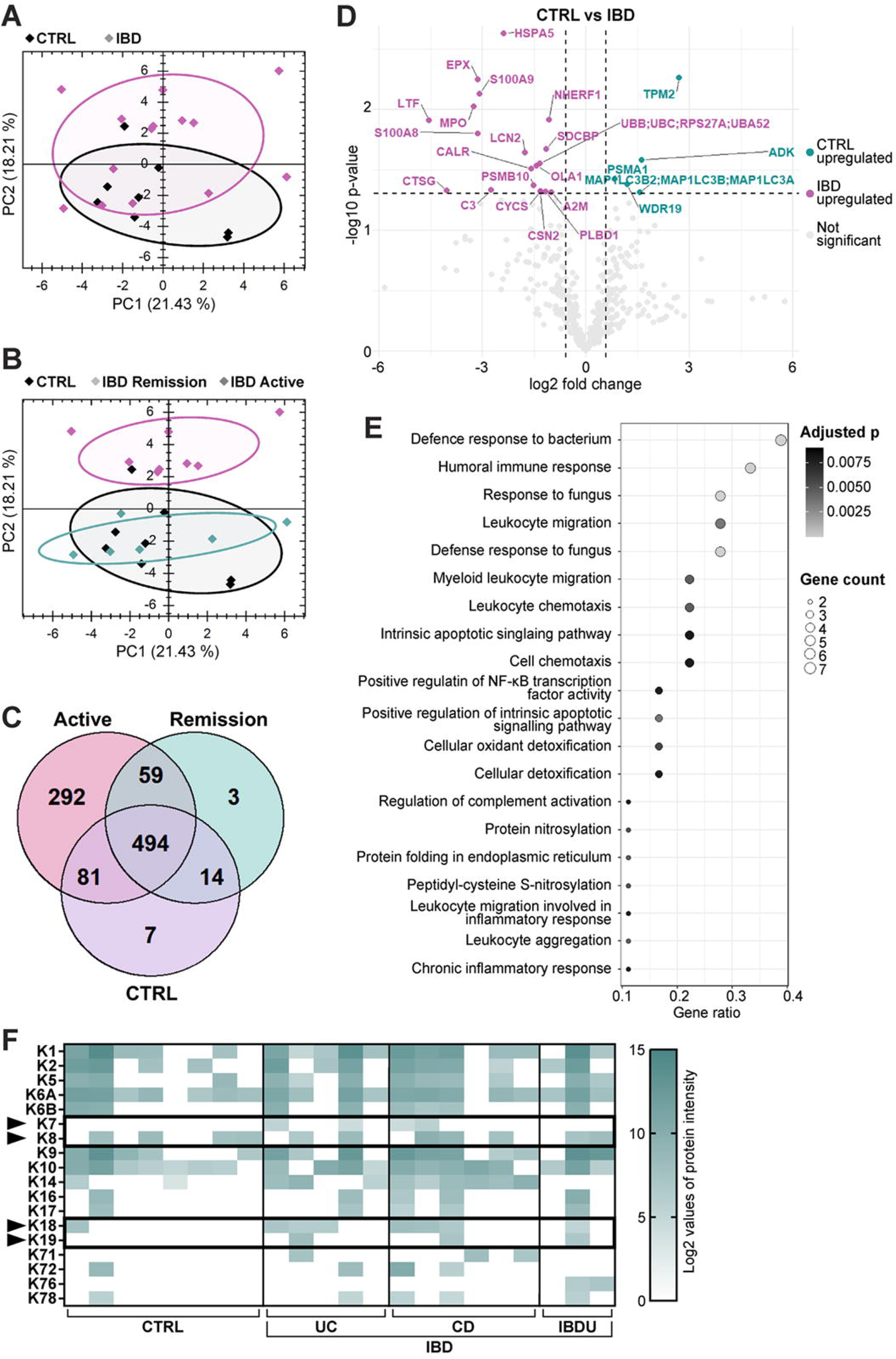
Proteomics analysis shows the effect of IBD and disease activity in the adolescent stool proteome. PCA plots presents clustering of IBD (dark gray, n= 14) and CTRL (black, n=9) stool samples (A), and comparison of IBD active (dark gray, n=8), IBD remission (light gray, n=6), and CTRL (black, n=9) stool samples (B). C) Venn diagram illustrates protein overlaps among stool samples from active IBD, IBD in remission, and CTRL groups. D) Volcano plot of differentially abundant proteins between IBD and CTRL stool samples. Each dot represents one protein. Labeled black or dark grey dots represent human proteins that were significantly different in abundance between groups and light gray points represent proteins that were not. Data was normalized across samples for consistent comparison. E) GO enrichment analysis of proteins more abundant in IBD stool. The top 20 GO BP categories are ranked by gene ratio. Bubble size reflects the number of genes involved in each term. F) Heatmap presenting the Log₂-transformed intensities of detected keratins in UC, CD, IBDU and CTRL stool samples. Intestinal keratins are highlighted.

Differential abundance analysis (CTRL vs IBD) revealed 23 proteins that fulfilled the predefined criteria (|log₂ fold change| ≥ 0.58, p ≤ 0.05). Among these, 18 proteins had higher and 5 proteins had lower abundance in IBD compared to CTRL (Figure 2D). Among the more abundant proteins in IBD were epithelial-associated proteins such as SDCBP, HSPA5 and NHERF1 (Figure 2D). GO BP overrepresentation analysis of the more abundant proteins in IBD yielded 65 enriched terms (p<0.05, BH-adjusted q<0.2) (Figure 2E). No enriched BP terms were detected among proteins less abundant in IBD.

Keratin proteins presence across sample groups was examined, independent of abundance criteria, which showed that K7 was detected in 4 samples, all from IBD patients. Fecal K19 was detected in IBD samples as well, whereas K8 and K18 were detected among both IBD and CTRL specimen (Figure 2F).

### Fecal K7 Protein Content Measured by Immunoassay can Differentiate IBD Patients from Non-IBD Patients

In order to investigate whether fecal K7 protein can be detected through immunoassay, K7 was immunoblotted from patient stool. K7 was found in >96% of IBD patient stool samples, where it was significantly higher than in non-IBD CTRLs, where K7 levels were close to or below the detection limit (Figure 3A-B). There were no significant differences in K7 levels between UC, CD and IBDU (Figure 3B). K7 protein was significantly elevated in the stool of patients with active IBD while stool samples from IBD patients in remission exhibited K7 levels on average comparable to those of non-IBD CTRLs (Figure 3C). The ROC curve of fecal K7 protein for distinguishing IBD fecal samples from non-IBD fecal samples yielded an AUC of 0.88 (95% CI) (Figure 3D). K7 values did not correlate with FC concentration (R^2^=0.04), (Figure 3E).

**Figure 3.**
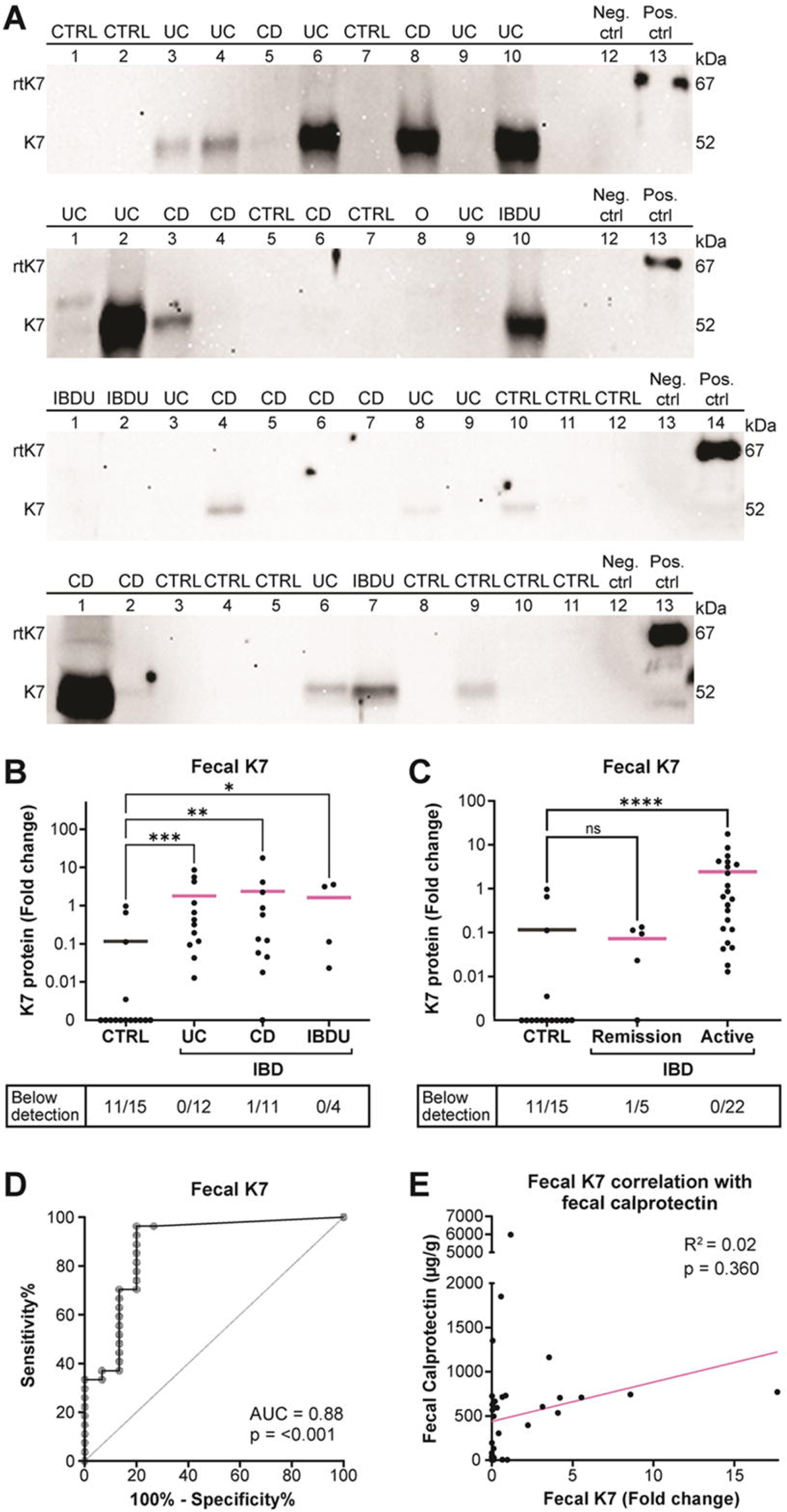
Elevated concentration of fecal K7 is observed in IBD patients with active disease. A) K7 protein expression levels measured in fecal sample lysates from CTRL (n=15), UC (n=12), CD (n=11), IBDU (n=4) and one non-IBD colitis (O) patients using Western blotting. Recombinant K7 (rtK7) was used as a positive control (Pos. ctrl), and yeast lysate as a negative control (Neg. ctrl). IBD samples stratified according to sub disease (B) and disease activity (C) were compared to CTRLs. Dots represent individual patients, and horizontal line indicate the median. D) ROC curves were generated for fecal K7 protein levels measured from patient stool samples by WB. E) Fecal K7 protein levels do not correlate with FC. Dots indicate individual patients. Statistical significance in B and C was determined using Kruskal-Wallis test and in D using simple linear regression analysis. *P<0.05, **P<0.01, and ***P<0.001.

### Fecal *KRT8* and *KRT18* Transcript Levels are Elevated in Active IBD

To determine whether there is an increase on mRNA level of keratin 7 (*KRT7)*, the stool samples were analyzed by qPCR. *KRT7* levels were low (Ct>36) and subject to major variation in all patient groups (Supplemental Figure 1A). A majority of the samples were below the detection limit. Thus, no significant differences in fecal *KRT7* levels between IBD subgroups or remissive/active IBD were observed (Supplemental Figure 1B). mRNA levels of the resident intestinal keratins (*KRT8*, *KRT18*, *KRT19* and *KRT20*) were, however, above detection limit in most of the fecal samples (Figure 4A-D), with *KRT8* and *KRT18* being significantly elevated in the IBD patient stool samples (Figure 4A-B). Fecal *KRT18* was furthermore significantly elevated in UC compared to CTRLs and in active disease compared to patients in remission (Supplemental Figure 2 and 3). ROC curve analysis to assess the diagnostic predictive value of *KRT8*, *KRT18*, *KRT19* and *KRT20* (Supplemental Figure 4) showed that *KRT18* exhibited the highest AUC of 0.78 (95% CI) (Supplemental Figure 4B). Pearson’s correlation analysis demonstrated that both *KRT18* and *KRT19* were significantly associated with increased fecal K7 protein levels (Figure 4E and Supplemental Figure 5). The strongest correlations detected were between fecal *KRT18* and *KRT19*, and fecal *KRT18* and *KRT20*.

**Figure 4.**
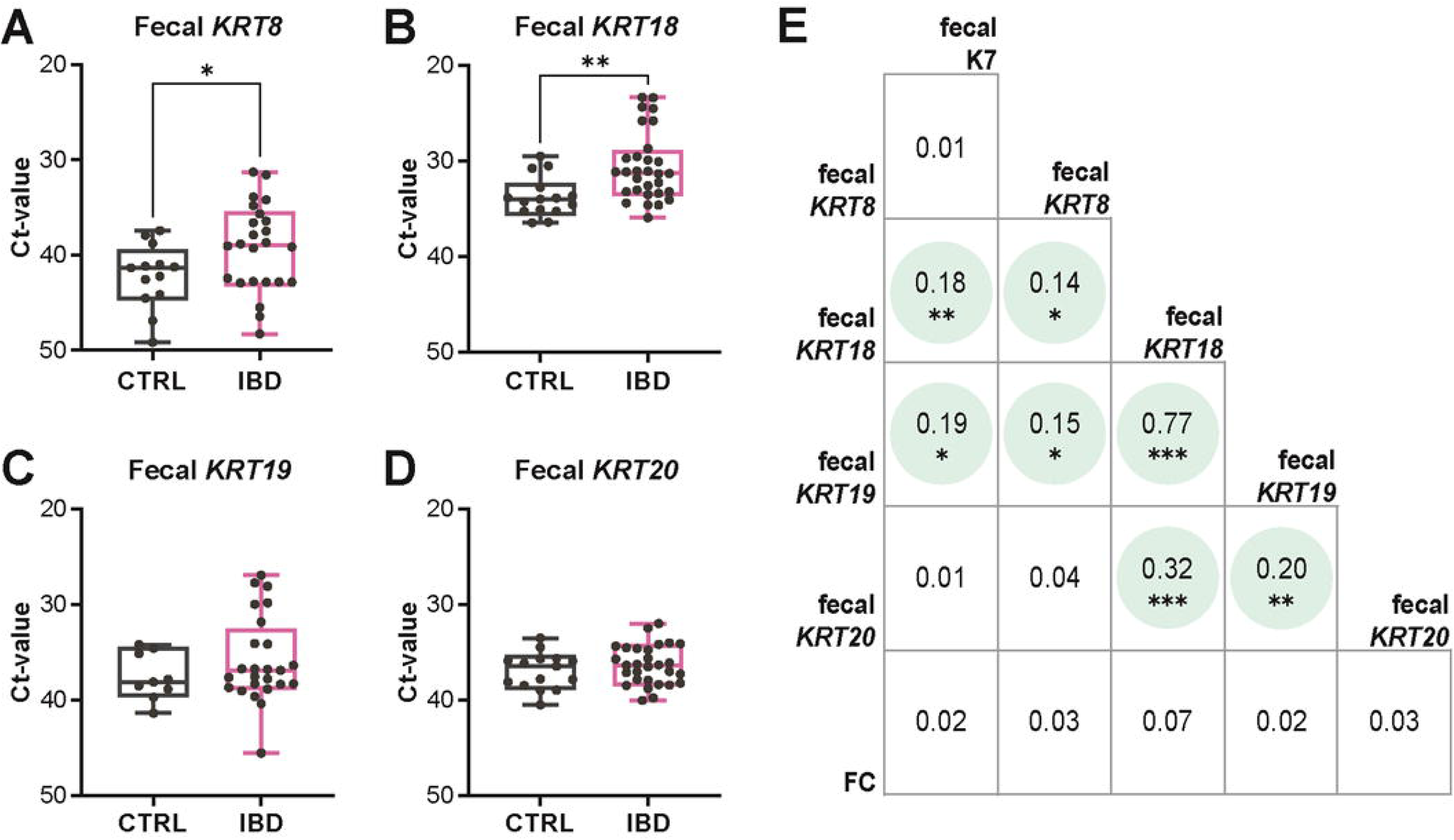
*KRT8 and KRT18* mRNA expression are increased in IBD patient stool samples. *KRT8* (A), *KRT18* (B), *KRT19* (C) and *KRT20* (D) mRNA expression levels measured in fecal sample lysates from CTRL (n=15) and IBD (n=27) patients using qPCR. The boxes extend from the 25th to the 75th percentiles, with the middle line representing median Ct-value and dots individual patients. Whiskers represent min-max values. E) Pearson’s correlation matrix for correlations between fecal (f) K7 protein levels, keratin Ct values, and FC. Values indicate R^2^, statistically significant correlations are marked in gray. Statistical significance comparing CTRL to IBD in A-D was determined using Kruskal-Wallis test. *P<0.05, **P<0.01, and ***P<0.001.

## Discussion

Our study is the first to demonstrate the feasibility of fecal K7 in noninvasively stratifying IBD patients from non-IBD patients. Colonocyte K7 expression was previously reported neo-expressed in adult IBD and in IBD-induced neoplasms, but not in microscopic colitis or irritable bowel syndrome [30,31,12,13,18], suggesting a high specificity for K7 in IBD. In our present study K7 was detected by immunoassay in the stool of nearly all adolescent IBD patients, resulting in a diagnostic performance with an AUC of 0.88. Notably, one patient with highly elevated FC was initially diagnosed with IBD but was later given another diagnosis. Interestingly, this patient was found to be negative for K7, further demonstrating K7 potential as a specific marker for IBD.

The mechanism behind intestinal K7 expression in IBD is yet to be defined. However, in accordance with previous results based on K7 levels assessed in IBD colon biopsies [12], fecal K7 and FC levels in IBD did not correlate, indicating that K7 is not a marker for acute inflammation. Keratins and other epithelia-derived markers could add a new dimension to IBD diagnostics in combination with immune cell activity assays, like the FC assay. Principally, fecal keratin could aid in stratifying IBD patients from patients with other intestinal diseases.

K7 was mainly increased in fecal samples from patients with active IBD, while patients in remission displayed levels closer to non-IBD. While the observed increase in stool keratins did not correlate with acute inflammation, the differences between patients in remission and with active IBD could point to some relation between chronic inflammation and keratin expression. Both increased epithelial shedding and excessive ulcerations in IBD can lead to the absence of keratin-expressing colonocytes, which makes it difficult to assess the actual keratin expression in inflamed colon epithelium [32]. A previous study focusing on intestinal biopsies indicated that keratin concentration varied individually as well as between disease subtypes [16].

It is yet unclear during which part of the disease progression K7 neo-expression occurs [12]. However, the differences between IBD patients in remission and active IBD patients, along with the large individual variations, would suggest that intestinal keratin expression and fecal keratin content change during different stages of the disease. This could partly be attributed to patients in remission having less epithelial damage, resulting in less erosion and a reduced amount of keratin expressing epithelial cells in stool. Evidence suggests that IBD could, in theory, be diagnosed years before the first symptoms, during a so-called pre-clinical period, but as of now there are no suitable specific biomarkers for this [2,33]. Considering that a disease phenotype with epithelial changes is already present in preclinical IBD [34–36], it would be of particular interest to determine at which point during the disease progression K7 is detectable in the tissue and stool.

The stool proteome exhibited a prominent inflammatory signature in IBD including neutrophil-derived proteins associated with the enrichment of inflammatory pathways, consistent with previous research on IBD fecal proteomics [37]. Additionally to these immune mediators, several epithelial-associated proteins including SDCBP [38], HSPA5 [39], and NHERF1 [40], were more abundant in IBD samples, likely reflecting epithelial stress and disruption of apical membrane integrity during mucosal injury. The elevated fecal levels of NHERF1 may indicate increased epithelial shedding into the intestinal lumen during active inflammation, which aligns with previous reports of NHERF1 epithelial downregulation in IBD [40].

Studies investigating potential fecal IBD biomarkers have mainly focused on immunological and microbial factors found in stool [41–43]. However, data on shed simple epithelial proteins has been lacking. Keratins in particular are considered common contaminants originating from sample handling in mass spectrometry, and therefore all keratins, including non-epidermal keratins such as those found in the intestinal epithelium, are often disregarded or even removed from proteomic data analysis as a part of contaminant libraries [44]. Although intestinal keratins were detected in our proteomics data, their inconsistent detection across samples limited their inclusion in differential abundance analysis. The wide dynamic range of stool proteomes tends to favor robust inflammatory proteins over epithelial proteins such as keratins, which may be less consistently detected in stool samples [45,46]. Overall, these findings suggest that stool proteomics capture inflammatory-associated proteins alongside epithelial derived components in IBD, although more sensitive approaches may be required for consistent quantification of epithelial proteins.

This study was primarily limited by the size of the cohort. Further studies with a larger cohort and a fully quantitative K7 assay will be required to validate a threshold value for K7 protein as a fecal biomarker. Due to the insoluble nature of keratins their measurement and analysis are easily impeded. Until now studies have commonly used harsh chemicals and detergents to solubilize keratins [47]. Likely due to these factors, compared to the proteomics, the immunoassay was more successful in identifying K7 by using harsher extraction conditions capable of isolating keratins from the stool matrix. These properties need to be overcome in order to develop fecal keratin quantification methods suitable for a hospital environment.

## Conclusion

Here we assessed the measurability of intestinal keratin protein and mRNA in stool samples and identified differences in fecal keratin prevalence between non-IBD, active IBD and IBD in remission. Fecal K7 in particular was shown to be feasible for noninvasive detection of IBD, and did not correlate with FC levels, suggesting potential complementary diagnostic value for K7 as a biomarker. Ultimately, these results support the development of keratin-based alternative noninvasive screening methods for IBD to complement current diagnostic methods.

## Author Contributions

Conceptualization: M.A.I., L.P., K-L.K., D.M.T.. Formal analysis: M.A.I., E.K., J.M.. Investigation: M.A.I., J.R., N.L., J.M.. Visualization: M.A.I., E.K.. Project administration: L.P., D.M.T.. Resources: A.N., R.V-H., O.K., K-L.K., D.M.T.. Supervision: O.K., K-L.K., L.P., D.M.T. Writing – original draft: M.A.I., E.K., L.P.. Writing – review and editing: All.

## Acknowledgements

We gratefully acknowledge the patients and their families for their contribution to the sample collection enabling this study. In addition, we thank all members of the Laboratory of Epithelial Biology at Åbo Akademi University for their assistance, especially Emelie Lassas for her help in experimental work. Mass spectrometry analysis was performed at the Turku Proteomics Facility, Turku Biosience Center, University of Turku and Åbo Akademi University. The facility is supported by Biocenter Finland.

## Data Availability Statement

All datasets generated and raw analyzed during this study are available from the corresponding author upon reasonable request. Relevant data supporting the findings of this study are included within the article and its supplementary materials. Proteomics raw data used in this study will be made publicly available upon publication, other relevant data supporting the findings of this study are included within the article and its supplementary materials. Shareable data excludes sensitive patient information containing potential personal identifiers, in accordance with applicable data protection regulations (Regulation (EU) 2016/679).

## Supplementary Tables

**Supplementary Table 1.** List of TaqMan probes.

| **Gene** | **Assay ID** |
| --- | --- |
| **name** | **(Thermo Fisher Scientific)** |
| *KRT7* | Hs00559840_m1 |
| *KRT8* | Hs01595539_g1 |
| *KRT18* | Hs02827483_g1 |
| *KRT19* | Hs00761767_s1 |
| *KRT20* | Hs00300643_m1 |

## Supplementary Figure Legends

**Supplemental Figure 1.**
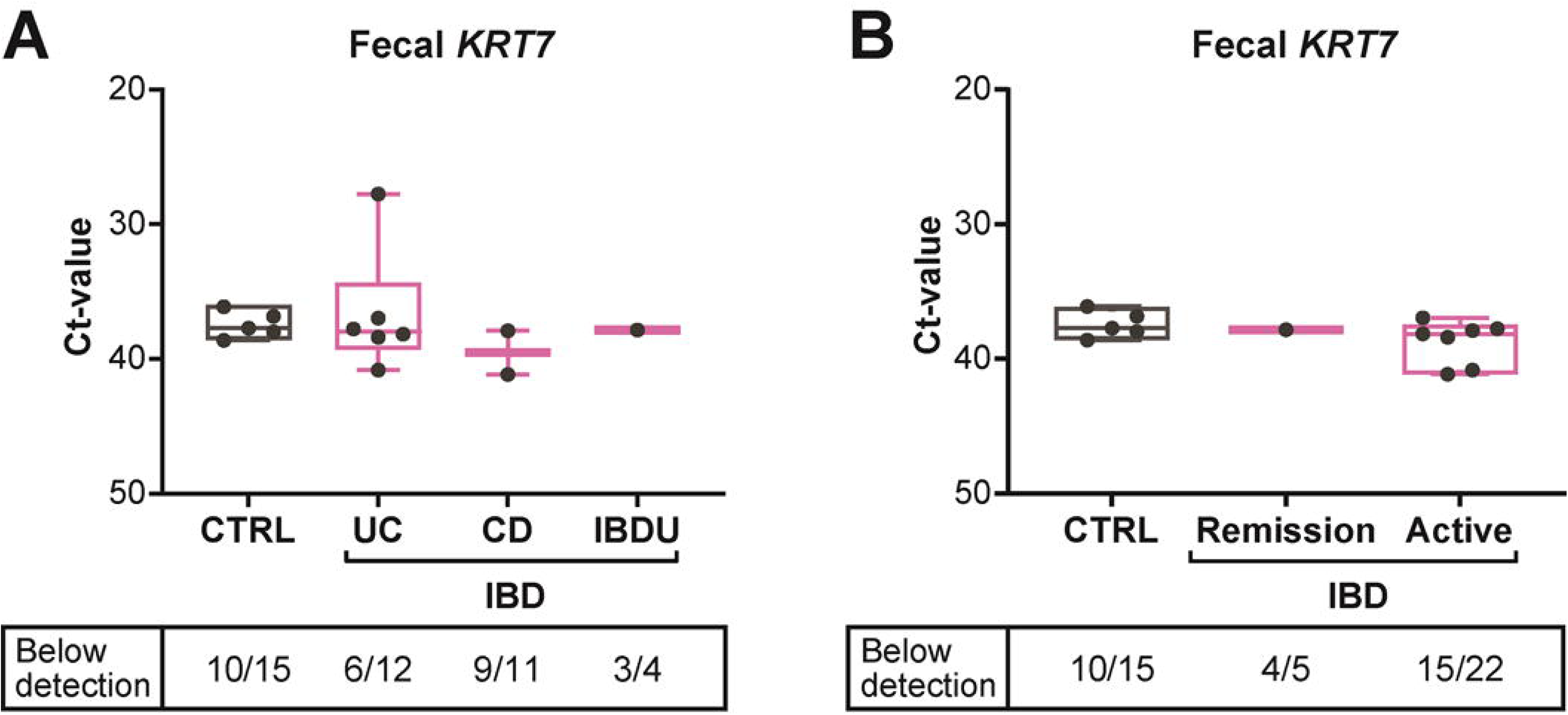
Fecal *KRT7* Ct values are mostly below detection limit, with no differences between IBD groups and CTRLs. A) *KRT7* expression measured in fecal sample lysates from CTRL (n=15), UC (n=12), CD (n=11) and IBDU (n=4) patients using qPCR. The boxes extend from the 25th to the 75th percentiles, with the middle line representing median Ct-value and dots individual patients. Whiskers represent min-max values. Statistical significances comparing IBD subgroups and activity to CTRL were determined using Kruskal-Wallis test.

**Supplemental Figure 2.**
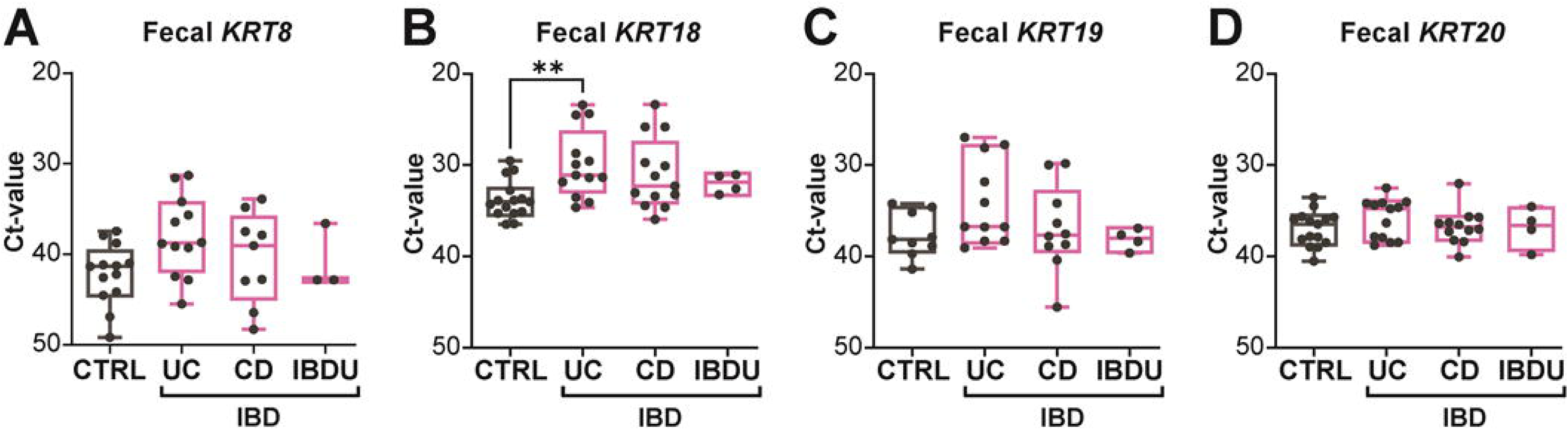
Fecal keratin mRNA levels in IBD subtypes. *KRT8* (A), *KRT18* (B), *KRT19* (C) and *KRT20* (D) mRNA Ct values were measured in fecal samples from CTRL (n=15), UC (n=12), CD (n=11) and IBDU (n=4) patients using qPCR. The boxes extend from the 25th to the 75th percentiles, with the middle line representing median Ct-value and dots individual patients. Whiskers represent min-max values. Statistical significance comparing IBD subgroups to CTRL were determined using Kruskal-Wallis test.

**Supplemental Figure 3.**
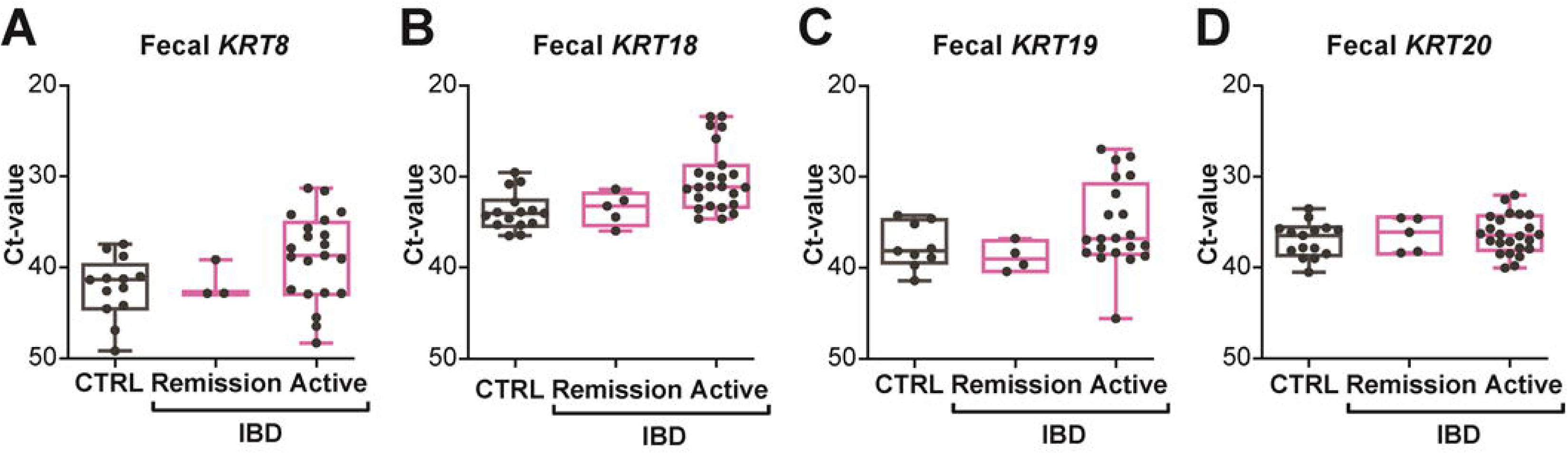
Fecal keratin mRNA levels in IBD according to disease activity. *KRT8* (A), *KRT18* (B), *KRT19* (C) and *KRT20* (D) mRNA Ct values were measured in fecal samples from CTRL (n=15) and IBD patients stratified according to disease activity (in remission, n=5, and active, n=22) using qPCR. The boxes extend from the 25th to the 75th percentiles, with the middle line representing median Ct-value and dots individual patients. Whiskers represent min-max values. Statistical significance comparing IBD activity to CTRL were determined using Kruskal-Wallis test.

**Supplemental Figure 4.**
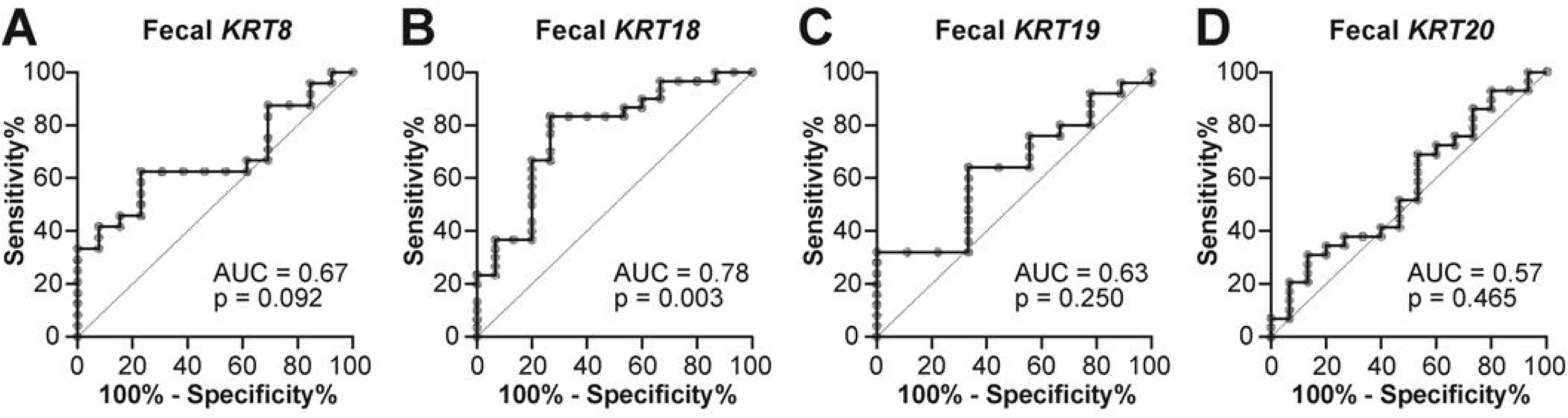
Evaluation of fecal keratin mRNA levels performance as IBD biomarkers. ROC curves were generated for fecal *KRT8* (A), *KRT18* (B), *KRT19* (C) and *KRT20* (D) Ct values.

**Supplemental Figure 5.**
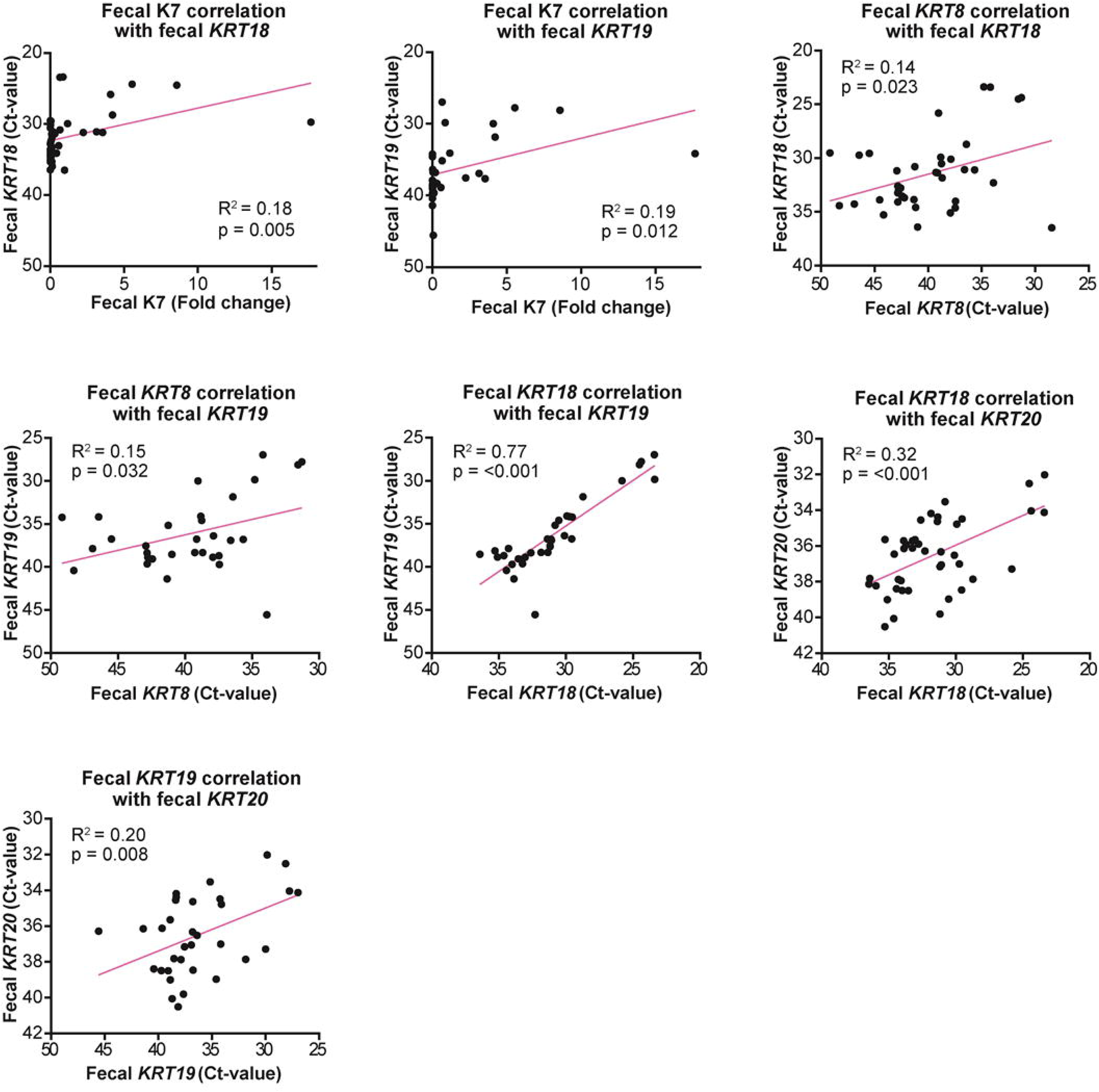
Linear correlation plots for all identified correlations between K7 protein and RNA levels of *KRT8, KRT18, KRT19*, and *KRT20* with statistical significance. Correlations are based on the data presented in Figure 4. Dots indicate individual patients. R^2^ and p-value were determined using Pearson’s simple linear regression analysis.

