## Supplemental material for "Keratin 7 protein presence in stool is indicative of active pediatric-onset inflammatory bowel disease"

**Supplemental Materials**

**Supplementary Table 1. List of TaqMan probes used**

| **Gene name** | **Assay ID (Thermo Fisher Scientific)** |
|---|---|
| *KRT7* | Hs00559840_m1 |
| *KRT8* | Hs01595539_g1 |
| *KRT18* | Hs02827483_g1 |
| *KRT19* | Hs00761767_s1 |
| *KRT20* | Hs00300643_m1 |

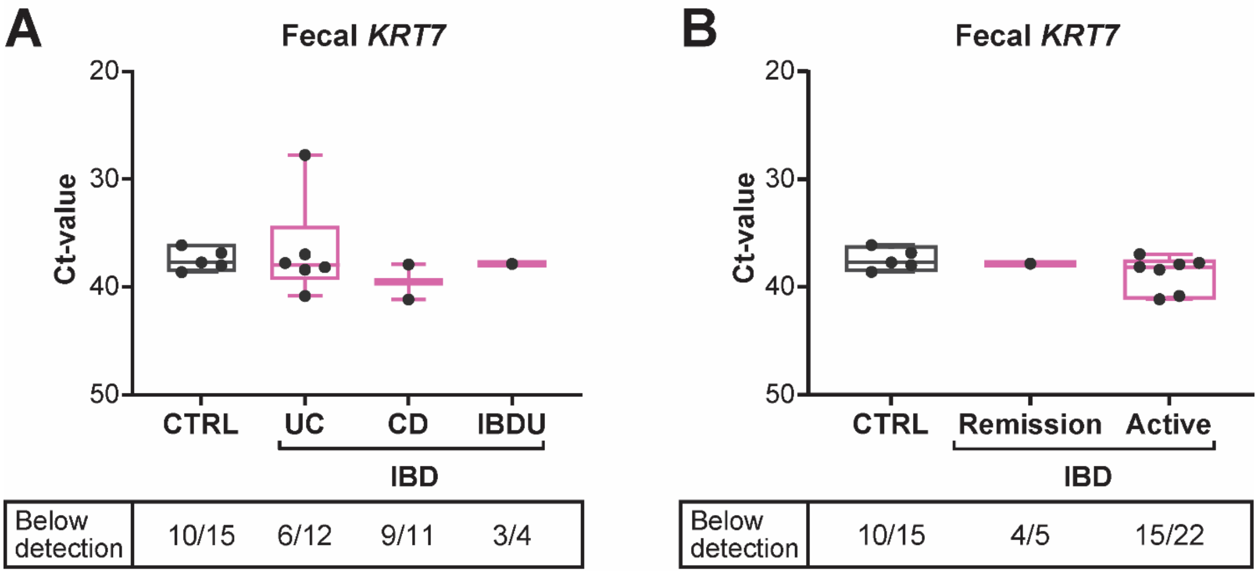


**Supplemental Figure 1. Fecal *KRT7* Ct values are mostly below detection limit, with no differences between IBD groups and CTRLs** A) *KRT7* expression measured in fecal sample lysates from CTRL (n=15), UC (n=12), CD (n=11) and IBDU (n=4) patients using qPCR. The boxes extend from the 25th to the 75th percentiles, with the middle line representing median Ct-value and dots individual patients. Whiskers represent min-max values. Statistical significances comparing IBD subgroups and activity to CTRL were determined using Kruskal-Wallis test.

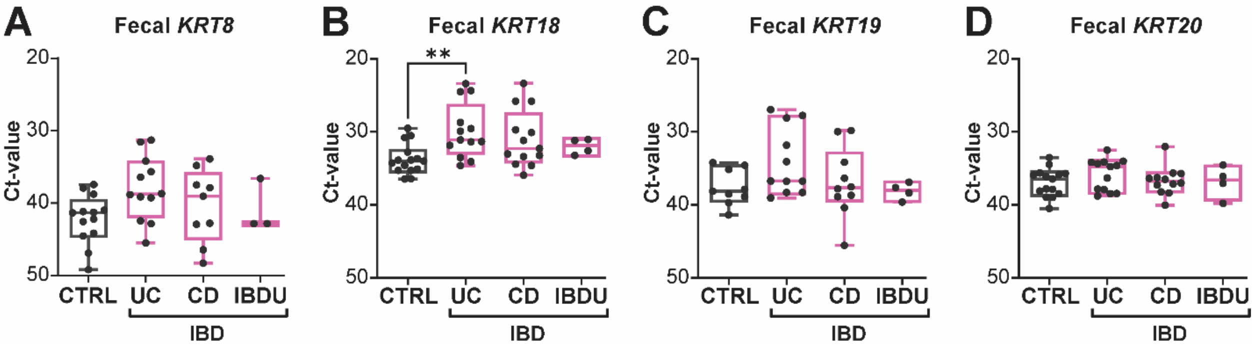


**Supplemental Figure 2. Fecal keratin mRNA levels in IBD subtypes.** *KRT8* (A), *KRT18* (B), *KRT19* (C) and *KRT20* (D) mRNA Ct values were measured in fecal samples from CTRL (n=15), UC (n=12), CD (n=11) and IBDU (n=4) patients using qPCR. The boxes extend from the 25th to the 75th percentiles, with the middle line representing median Ct-value and dots individual patients. Whiskers represent min-max values. Statistical significance comparing IBD subgroups to CTRL were determined using Kruskal-Wallis test.

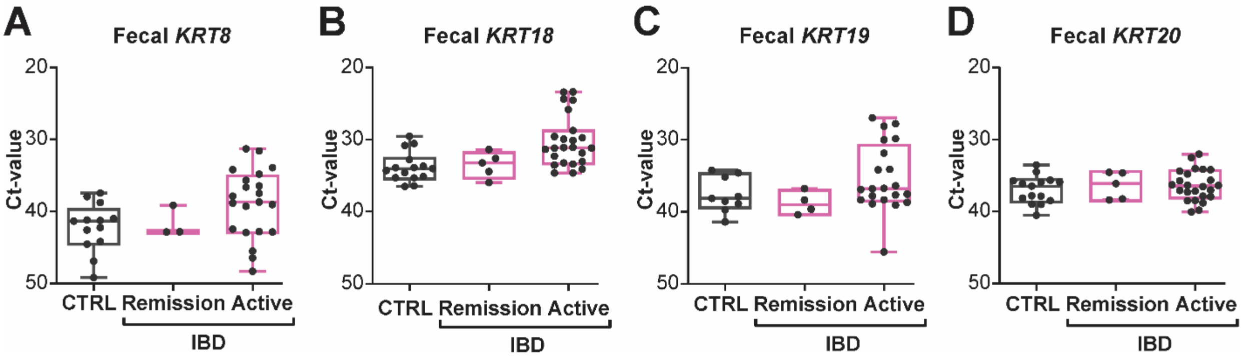


**Supplemental Figure 3. Fecal keratin mRNA levels in IBD according to disease activity.** *KRT8* (A), *KRT18* (B), *KRT19* (C) and *KRT20* (D) mRNA Ct values were measured in fecal samples from CTRL (n=15) and IBD patients stratified according to disease activity (in remission, n=5, and active, n=22) using qPCR. The boxes extend from the 25th to the 75th percentiles, with the middle line representing median Ct-value and dots individual patients. Whiskers represent min-max values. Statistical significance comparing IBD activity to CTRL were determined using Kruskal-Wallis test.

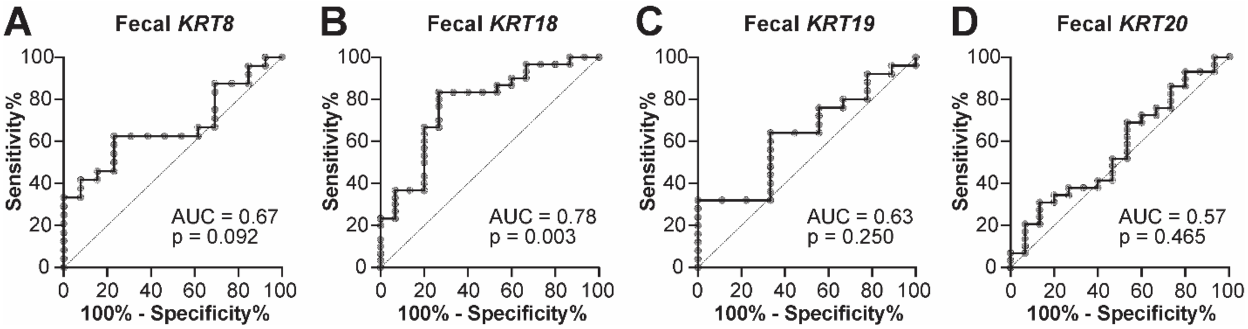


**Supplemental Figure 4. Evaluation of fecal keratin mRNA levels performance as IBD biomarkers.** ROC curves were generated for fecal *KRT8* (A), *KRT18* (B), *KRT19* (C) and *KRT20* (D) Ct values.

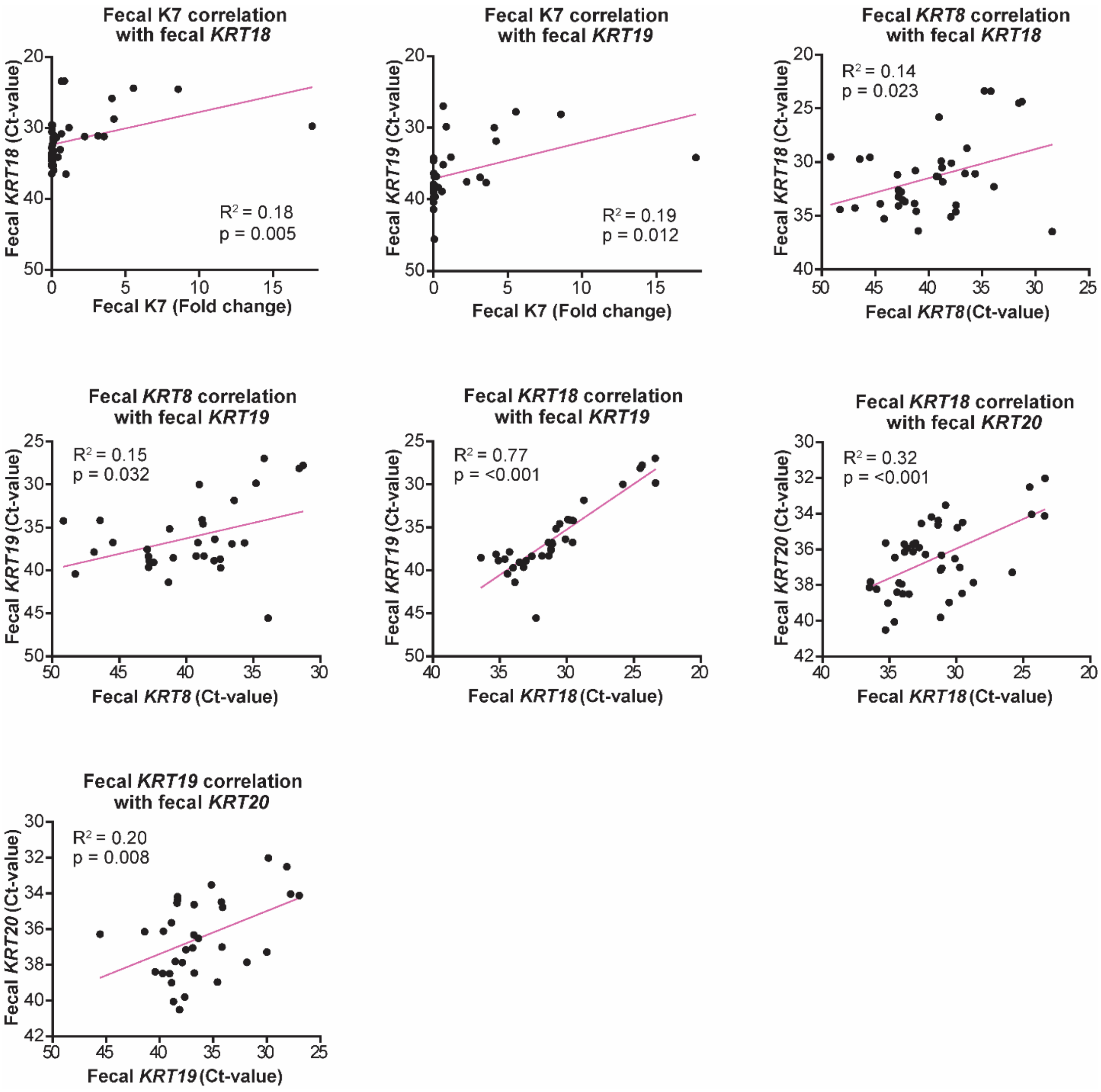


**Supplemental Figure 5. Linear correlation plots for all identified correlations between K7 protein and RNA levels of *KRT8, KRT18, KRT19*, and *KRT20* with statistical significance**. Correlations are based on the data presented in figure 4. Dots indicate individual patients. $R^2$ and p-value were determined using Pearson's simple linear regression analysis.
